# Raloxifene reduces sex- and age-related intervertebral disc degeneration in mice by estrogen signaling

**DOI:** 10.1101/2021.06.29.449482

**Authors:** Neharika Bhadouria, Paul Niziolek, Omar El Jordi, Alycia G. Berman, David McKinzie, Joseph M. Wallace, Nilsson Holguin

**Author notes:** Nilsson Holguin, Ph.D., Assistant Professor, Department of Mechanical and Energy Engineering, IUPUI, 723 W. Michigan St., Indiana Center of Musculoskeletal Health, IUPUI, 635 Barnhill Drive. Indianapolis, IN, 46202, USA, Voice.

## Abstract

Estrogen agonist raloxifene is an FDA-approved treatment for osteoporosis in postmenopausal women that may also be a promising prophylactic for painful intervertebral disc (IVD) degeneration. Here, we hypothesized that raloxifene would augment IVD structure and reduce neurokinin-1 (substance P) in young and old mice by stimulating estrogen signaling. 2.5 month (male and female) and 22.5 month (female) C57Bl/6J mice were subcutaneously injected with raloxifene hydrochloride (5x/week, 6week, n=7-9/grp). Next, to determine the impact of estrogen-deficiency to IVD structure and substance P, female mice were ovariectomized (OVX) at 4mo and tissues from OVX and sham-operated mice were harvested at 6mo (n=5-6/grp). First, compared to male IVD, female IVD expressed less *col2* and *osterix* transcription, early markers of IVD degeneration. Irrespective of sex, raloxifene increased the transcriptional expression for extracellular matrix anabolism, proliferation, notochordal cells (vs chondrocyte-like cells) and estrogen signaling in young IVD. Next, we determined that biological sex and aging each induced structural features of lumbar IVD degeneration. Therapeutically, injection of raloxifene countered these features by increasing IVD height in young mice, preventing mild sex-related IVD degeneration in young female mice and partially reversing age-related IVD degeneration in old female mice. Further, estrogen agonist raloxifene upregulated er-α protein and downregulated substance P protein in young and old IVD. By contrast, estrogen-deficiency by OVX increased IVD degeneration and substance P protein in IVD cells. Similarly, substance P protein in vertebral osteocytes was upregulated in females relative to males and by estrogen-deficiency and downregulated by raloxifene. Overall, raloxifene augmented IVD structure and reduced substance P expression in young and old female murine IVD, whereas estrogen-deficiency increased substance P in the spine. These data suggest that raloxifene may potentially relieve painful IVD degeneration in postmenopausal women induced by biological sex, estrogen-deficiency and advanced age.

Graphical Abstract
Injection of raloxifene promotes IVD health by engaging estrogen and Wnt signaling to promote cell proliferation and IVD structure. Differential estrogen signaling by raloxifene and ovariectomy regulated nerve signaling protein substance P in the spine. Raloxifene may also bind water to collagen to promote hydration. Acan: aggrecan, AF: annulus fibrosus, NC: notochordal cell, NP: nucleus pulposus

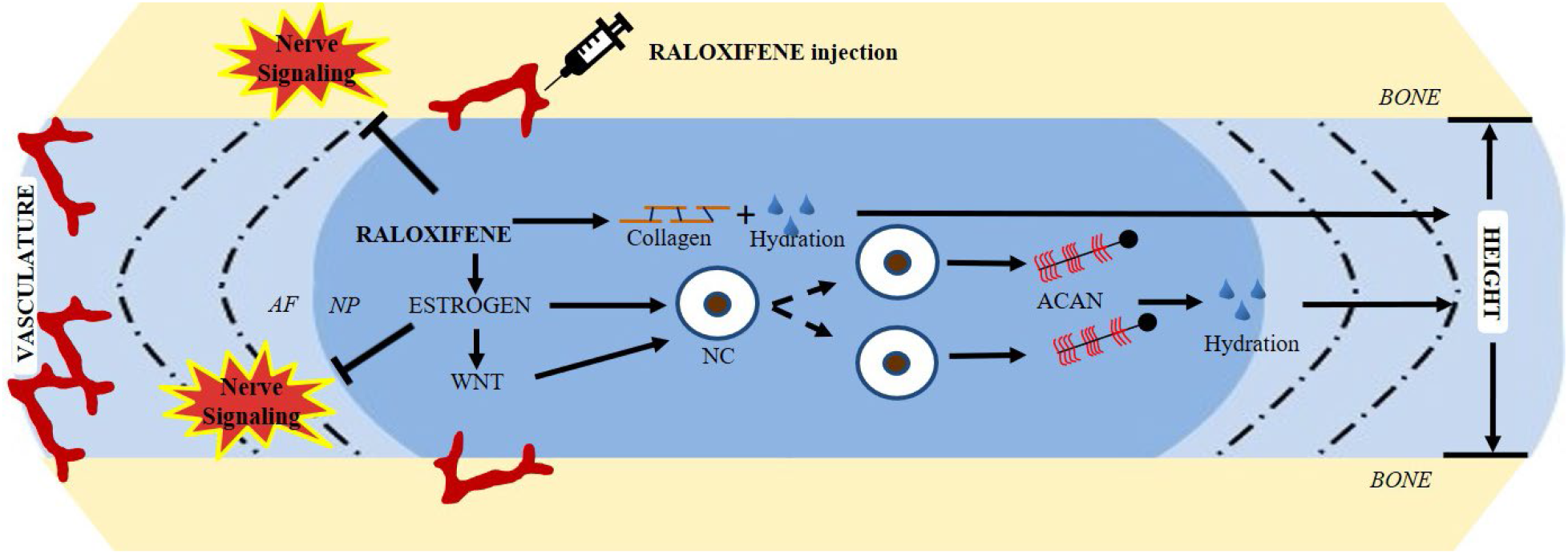

## Introduction

Currently, there are no FDA-approved pharmacological treatments to promote intervertebral disc (IVD) structure or prevent IVD degeneration^(1,2)^, a major contributing factor of low back pain^(3,4)^. 1 in 4 individuals suffer from chronic low back pain and the costs from healthcare treatment and lost wages are estimated to be in the hundreds of billions of dollars in the United States^(5)^. IVD degeneration is characterized by IVD height loss, extracellular matrix (ECM) breakdown, dehydration of the central nucleus pulposus and cell loss^(6–9)^. The hypo-vascularity and hypo-cellularity of the IVD challenge its self-repair and access by regenerative therapies^(10)^. These intrinsic limitations have motivated the development of promising approaches to replace severely degenerated IVDs with tissue-engineered ones^(11)^, but these invasive approaches preclude the prevention of native IVD degeneration and are not designed to spare patients the subsequent pain or cost of surgery. While mechanical stimuli can protect the IVD^(12–14)^, a pharmacological may be more consistent regenerative therapy. Pharmacological agents administered systemically can reach the IVD^(15)^ and, considering the potential relationship between osteoporosis and IVD degeneration^(16)^, treatments for bone structure may also target regenerative pathways in the IVD.

Pre- and post-menopausal women develop greater IVD degeneration than age-matched men^(17,18)^ and experience pain more frequently and at higher intensities^(19,20)^. Raloxifene-use in women relieves self-reported back pain ^(21)^and increases IVD height^(22)^ for unclear reasons. Raloxifene hydrochloride is an FDA-approved non-uterine-targeting selective estrogen receptor modulator (SERM) that suppresses bone resorption by binding to estrogen receptors (er) in osteoclasts^(23,24)^. The dramatic 50% reduction of vertebral fracture, which was previously unexplained by the mild 4% increase in bone mineral density^(25)^, may also be associated with biophysical binding of water to collagen^(26)^ than er signaling in bone^(27)^. Therefore, Raloxifene may be beneficial to the IVD by increasing water content and at least 2 other mechanisms:

i. Raloxifene may engage Wnt signaling to promote the maintenance of notochordal cells and ECM of the IVD. The nucleus pulposus is the hydration core of the IVD and healthy IVD are mostly populated by notochordal cells, rather than chondrocyte-like cells. Notochordal cells require Wnt signaling to maintain their cell phenotype^(28)^ and are better equipped than chondrocyte-like cells to produce ECM^(29)^. Aging and IVD degeneration^(30,31)^ perpetuate ECM degradation^(32,33)^ by reducing Wnt signaling, which triggers the replacement of notochordal cells by chondrocyte-like cells. Contrarily, stabilization of Wnt signaling transcription factor *β-catenin* in the nucleus pulposus promotes notochordal cell proliferation, augments ECM anabolism and protects the IVD from injury-induced ECM degradation^(33)^. Therefore, raloxifene promotes the proliferation of IVD cells^(34)^ by binding to er-α^(30)^ and potentiating Wnt/β-catenin signaling^(31,35)^.
ii. Raloxifene may regulate pain intensity via nociceptive processing in the central neural system^(36)^. Discogenic pain by substance P (neurokinin-1) has long been associated with IVD degeneration^(37)^. Substance P (SP) is a nerve signaling neurotransmitter and nociceptive pain marker that is negatively associated with er-α protein expression in the IVD of postmenopausal women^(38)^. Considering that raloxifene binds Erα, raloxifene may in turn reduce substance P. Here, we determined whether systemic delivery of estrogen-agonist raloxifene and estrogen-deficiency altered IVD structure, the cellular expression of substance P, ECM markers and notochordal cell-related markers. Systemic drug delivery avoids iatrogenic IVD degeneration by local injection, but the effective quantity is dramatically reduced at the IVD because of its limited vascularization. Therefore, the pharmacologic must be potent, yet safe for repeated delivery. The FDA approved raloxifene for daily-use in 1997 and it does not negatively engender back pain^(29,36)^. In order to test the impact of repeated-use, we injected raloxifene to C57Bl/6J mice 5x/week for 6 weeks. In our hands, repeated injection of raloxifene improved the structural and cellular composition of young-adult IVD and reduced the expression of pain-related neuropeptide substance P. In old mice, raloxifene similarly reduced substance P and partially improved IVD structure. In ovariectomized mice, estrogen-deficiency increased substance P and induced mild IVD degeneration showing the relation between estrogen and pain biomarker.

## Materials and Methods

### Mice

The objectives of this study were to investigate the effects of systemic delivery of raloxifene in young and old IVDs. We further clarified the role of estrogen in IVD by ovariectomy in mice. This in-vivo study was approved by Indiana University of School of Medicine Laboratory Animal Resource Center, Institutional Animal Care and Use Committee (IACUC). Tissues of the same set of mice were used in two different studies with different protocols. Mice were housed in a 12-hour light/dark cycle and fed standard chow. In this study, 2.5-month-old CON male and female (C57Bl/6J, n=7-9/sex/group) were injected with 0.5 mg/kg of raloxifene hydrochloride (SIGMA) for 6 weeks, 5x/week (Ral) and harvested at 4 month of age. Next, 22.5 month old female mice purchased from Jax (C57Bl/6J, n=8/group) were injected with the same dose and frequency of raloxifene hydrochloride or vehicle PBS (VEH) and harvested at 24-month of age. Lastly, 4-month old female mice (C57Bl/6J, n=4-5/group) were ovariectomized and harvested at 6-month of age (OVX). Control mice were sham-operated (SHAM). Mice were euthanized by hypoxia as a primary means and cervical dislocation as a secondary means. Lumbar and tails were harvested, and the spinal segments were divided-up for specific testing (Table S1).

### Histology and Immunohistochemistry

All IVDs were run in a single batch. Motion segments were fixed in 15 mL of 10% formalin on a rocker for 24 hours, submerged in 70% ethanol, embedded in paraffin, and sectioned (5 μm). In short, the nucleus pulposus (NP), annulus fibrosus (AF), and boundary between the two structures were scored based on structural properties. Then, added for a total IVD score between 0-14, with greater scores denoting IVD degeneration. Ki67 positive cells were counted within 200 μm of length from the NP boundary (20x image). This region of interest was chosen as ki-67 expressing cells were dominating in this region for all samples. For quantification of other protein expressions, the whole of NP or AF were considered as regions of interest for cell count. For β-catenin (NP), the total number of positively stained cells in the NP were counted and compared. Histological images of IVDs were used to measure IVD structures and were measured using ImageJ to determine if there were any changes.

### Magnetic Resonance Imaging (MRI)

Motion segment CC6-7 was submerged and wrapped in 1x PBS-soaked gauze overnight until imaged. T2 weighted imaging was completed on the Bruker BioSpin 9.4 T MRI, using a 0.4 mm slice thickness. Two samples were stacked in a glass tube to remain upright, and two glass tubes were placed, separated by foam composite, inside of a 15mL tube to ensure samples would not move while being imaged.

### Micro-computed tomography

Motion segments L6-S1 and CC6-7 were harvested and submerged in 1x PBS prior to imaging. Specimens were imaged using the Bruker SkyScan 1272 Micro-CT at a resolution of 8 micrometers. Images of the motion segment were contoured around the periosteal and the endosteal of the bone. For the trabecular analysis, the growth plate was used as a landmark and trabecular bone analysis consisted of the next 30 consecutive images^(39,40)^. For cortical analysis, the longitudinal center of the bone was identified, and 15 images above and below were analyzed using the Bruker CTan64 microCT software. Parameters measured include bone volume fraction (BV/TV), trabecular number (Tb.N), trabecular thickness (Tb.Th) for trabecular bone and cross-sectional thickness for cortical bone, using a lower threshold of 60 and upper threshold of 225 for analysis.

### QPCR

L3-5 IVDs and CC8-10 IVDs were harvested, frozen in liquid nitrogen, pulverized and suspended in TRIZOL (#15596018, Invitrogen) until further processing^(41)^. RNA isolation and purification steps were followed (RNeasy mini kit, Qiagen) and RNA concentration was quantified (Nanodrop). CDNA was synthesized (iScript, BioRad) from 400 ng of total RNA for the following Taqman probes (Life Technologies); *Acan* (Mm00565794_m1), *β-catenin* (Mm01350387_g1), *Col1a1* (Mm00801666_g1), *Col2a1* (Mm01309565_m1), *Er-α* (Mm00433149_m1), *Foxa2* (Mm01976556_s1), *Hsp70* (Mm01159846_s1), *Krt19* (Mm00492980_m1), *Osx* (Mm04209856_m1), *Runx2* (Mm00501584_m1), *Mmp3*(Mm00440295_m1), *Lef1* (Mm00550265_m1), *Rela* (Mm00501346_m1) and *Pi3k* (Mm01282781_m1). Relative gene expression was normalized to *18s* (Hs99999901_s1) for each group and then experimental values were normalized to the average of the CON value (2^−ΔΔCT^).

### Analysis

IVD morphology and histological score were determined from Safranin-O/Fast green images. Degeneration scoring was the average of 5 independent observers^(20,42)^. Methyl green staining was the counterstain for the IHC staining of β-catenin (#9562S, Cell Signaling), ki-67 (# PA5-19462, Invitrogen), er-α (# MA3-310, Invitrogen) and substance P (ab14184, Abcam). For cell quantification in the NP or AF, the image was thresholded for intensity to demarcate the positive cell by semi-automation using ImageJ (NIH). Quantitative PCR was accomplished as previously described^(38)^. Briefly, relative gene expression was normalized to *18s* (Hs99999901_s1) for each sample and then experimental values were normalized to the CON average CT value (2^−ΔΔCT^).

For MRI, the motion segment was imaged in a sagittal orientation using a 0.052 x 0.052mm voxel resolution in the x-y direction and a 0.4 mm voxel resolution in the z-direction (16 averages/slice). Water area and intensity of the IVD were determined and multiplied to estimate the hydration content of the IVD (MRI Index) using ImageJ (NIH). A Bruker SkyScan 1172 MicroCT imaged functional spinal units at a resolution of 8 μm and was used to quantify vertebral cancellous and cortical bone structure.

### Statistics

Statistical analyses were performed using Statistical Package for the Social Sciences (SPSS Version26) software. A two-way analysis of variance (ANOVA) was used to compare the RAL to CON and determine the main effects of treatment, sex and their potential interaction in lumbar and tail IVDs. In case of significant interaction, Tukey post hoc testing was further conducted. A one-way ANOVA was used to compare the effect of aging in female control IVD. A Student’s t-test compared VEH to RAL groups, SHAM to OVX and the protein expression of β-catenin between CON and RAL groups. A Chi-squared test was used for NP cell band incidence measured by MRI. Data are represented as box plots with mean (marked as cross (x) and maximum/minimum whiskers or mean ± standard deviation. A p-value of <0.05 was considered statistically significant.

## Results

### Raloxifene upregulated pro-anabolic gene expression in male and female lumbar IVDs

Aggrecan (*Acan*), collagen II (*Col2a1*) and collagen I (*Col1a1*) are key components of the ECM of the IVD. Between sexes, the gene expression suggests that female IVDs were more degenerated than male IVDs. Compared to male IVDs, female IVDs expressed less *col2a1* and *osx* (osterix) gene expression and more *rela* (downstream of estrogen signaling) and *hsp70* (cellular stress marker) gene expression (Fig. 1, S1). Therapeutically, raloxifene treatment potently increased gene expression of *acan* by 4-5-fold and *col2a1*/*1a1* by 2-3-fold in IVDs. Cellular expression for notochordal and chondrocyte-like cell markers supported the anabolic gene expression induced by raloxifene. Systemic injection of raloxifene increased the gene expression for notochordal cell transcription factors, *foxa2* and *krt19* by 1-4-fold, while not altering the gene expression of chondrocyte-like cell transcription factors *osx* and *runx2*.

**Fig. 1.**
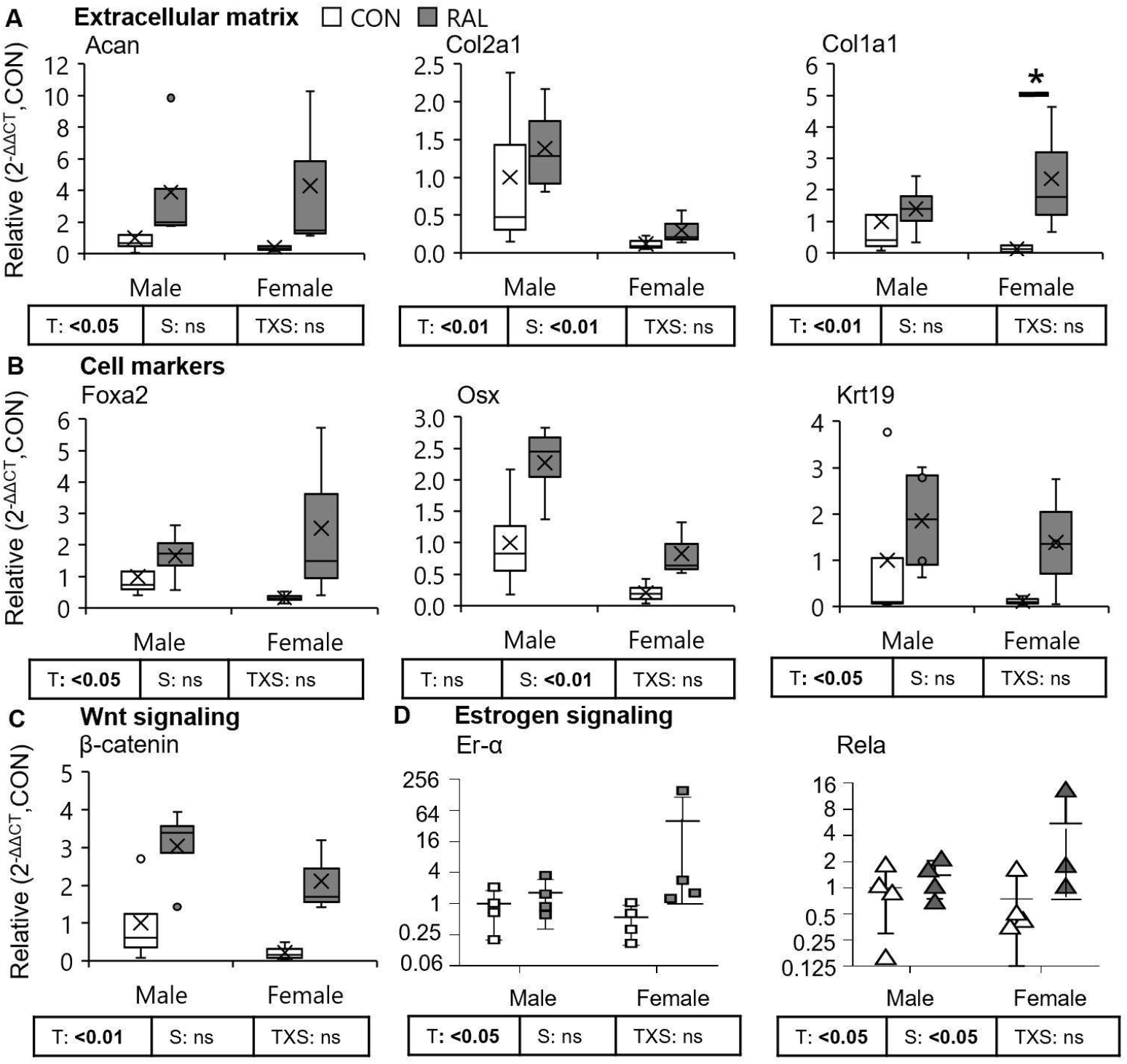
Raloxifene upregulated pro-anabolic gene expression in young male and female lumbar IVDs. Gene expression analysis for (A) extracellular matrix (*Acan*, *Col2a1 and Col1a1*), (B) transcription factors associated with cell type (*Foxa2*, *Osx, Krt19*), (C) Wnt signaling (*β-catenin*), and (D) estrogen signaling (*Er-α*, n=4-5/sex/group). Data are represented as box plots with mean marked as cross (x), 25/75% deviation lines and maximum/minimum whiskers. Control (CON) vs Raloxifene-treated (RAL), T: CON vs RAL, S: Male vs Female, TxS: Interaction. P<0.05. Scale bar: 100 μm.

### Raloxifene stimulated nucleus pulposus cell proliferation and canonical Wnt signaling

Concomitant to ECM anabolism, raloxifene stimulated proliferation pathways Wnt signaling and estrogen signaling. Specifically, *β-catenin* gene expression of male and female IVDs were upregulated by raloxifene. Canonical Wnt/β-catenin signaling is a known regulator of cell fate and proliferation^(43)^. Therefore, we determined the proliferation of nucleus pulposus cells using staining for ki67 (Fig. S2A). Raloxifene increased the number of proliferating cells (arrow) in nucleus pulposus by 135% and 67% in males and females, respectively (Fig. S2A’), but males were more responsive. Similarly, injection of raloxifene increased the number of total cells by 49% (Fig. S2A’’). Corroborative of the gene expression for *β-catenin* and pattern of proliferation in the nucleus pulposus, raloxifene increased the protein expression of β-catenin in the nucleus pulposus of female mice by 62% (Fig. S2B, B’). No annulus fibrosus cells were noted to express β-catenin protein.

### Raloxifene prevented sex- and age-related lumbar IVD degeneration

Compared to male control IVDs at 16 weeks of age, female control lumbar IVDs were more degenerated by 74%, based on histological scoring (Fig. 2A, A’). By contrast, injection of raloxifene for 6 weeks reduced the lumbar IVD degeneration score by 41% (Fig.2A’) in young male and female IVDs. Loss of IVD height is a common feature of IVD degeneration^(44,45)^. Irrespective of biological sex, injection of raloxifene increased IVD height by 11% (Fig. 2A’’) and nucleus pulposus area by 14% (Fig. S3). With respect to aging and compared to young female IVDs, old female IVDs were degenerated by 146% compared to the vehicle group, based on IVD scoring (Fig. 2B, B’). Raloxifene reduced the age-related IVD degeneration score by 57% with no change in IVD height (data not shown).

**Fig. 2.**
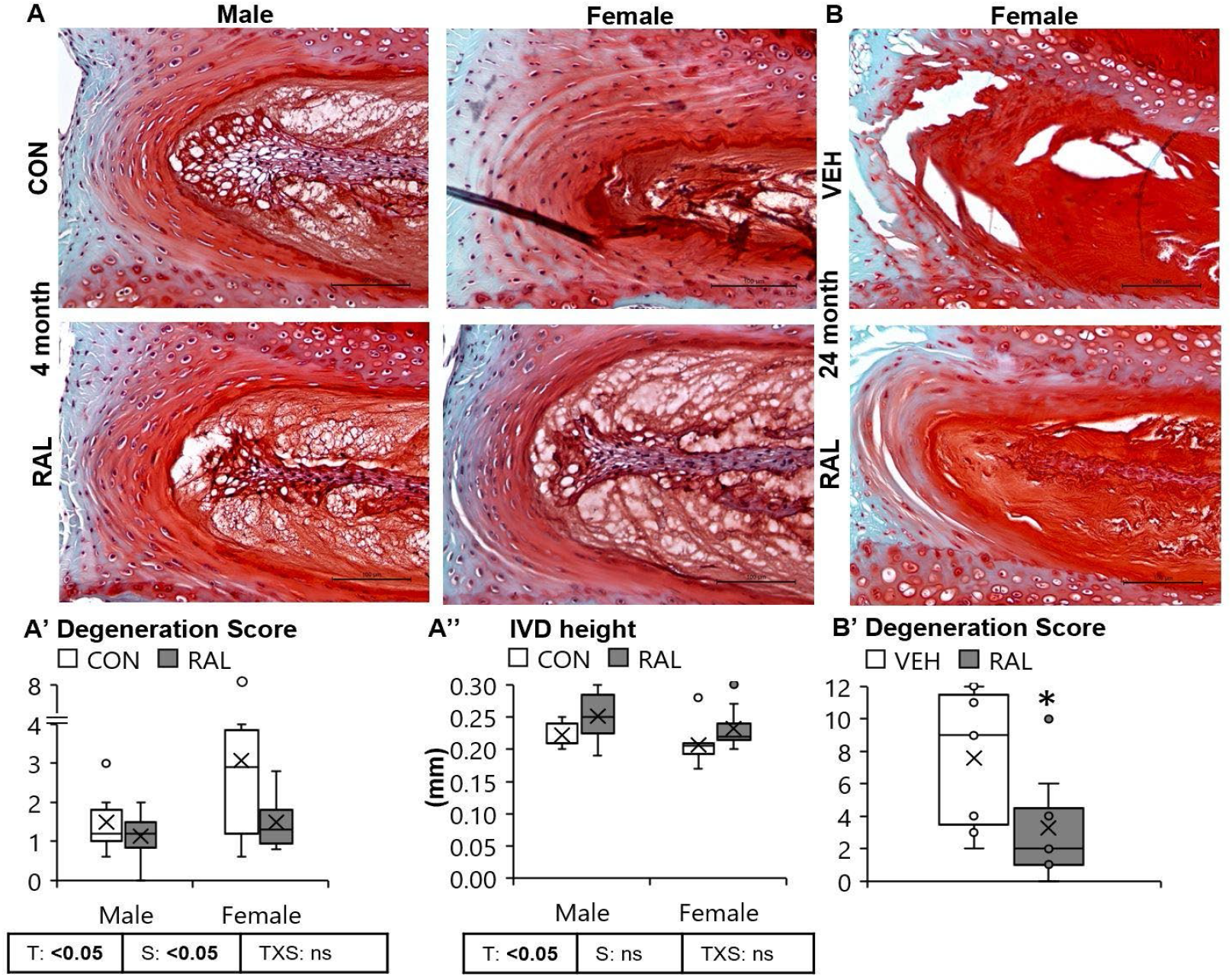
Raloxifene improved lumbar IVD degeneration score and IVD morphology. (A) Evaluated using Safranin-O/Fast Green staining, raloxifene (A’) reduced the IVD degeneration score and (A’’) increased IVD height of male and female young mice. (B’) Similarly, raloxifene reduced theIVD degeneration score of old female IVDs. Data are represented as box plots with mean marked as cross (x), 25/75% deviation lines and maximum/minimum whiskers. Control (CON, n=8-9/sex/group) vs Raloxifene-treated (RAL), T: CON vs RAL, S: Male vs Female, TxS: Interaction, Vehicle (VEH, n=8/group) vs Raloxifene Treated (RAL), *: VEH vs RAL (Female), p<0.05. AF: annulus fibrosus, NP: nucleus pulposus. Scale bar:100 μm.

### Raloxifene treatment stimulated estrogen signaling and reduced discogenic Substance P in young and old IVDs

Substance P is a nerve signaling neurotransmitter and nociceptive pain marker that is negatively associated with er-α protein expression in the IVD of postmenopausal women^(38)^. Considering that raloxifene binds er-α, raloxifene may in turn reduce substance P. Unlike the nucleus pulposus-only expression of β-catenin, raloxifene increased the er-α protein expression in the nucleus pulposus and annulus fibrosus (Fig. 3A-A’’), and the pattern of the *er-α* gene expression for whole IVD reflected the protein expression in the annulus fibrosus in young IVDs (Fig. 1D). Compared to young male IVDs, young female IVDs were more responsive to raloxifene treatment with a 2-fold increase in er-α-positive annulus fibrosus cells (Fig. S4A, B). 24-mo old female IVDs responded similarly to treatment.

**Fig. 3.**
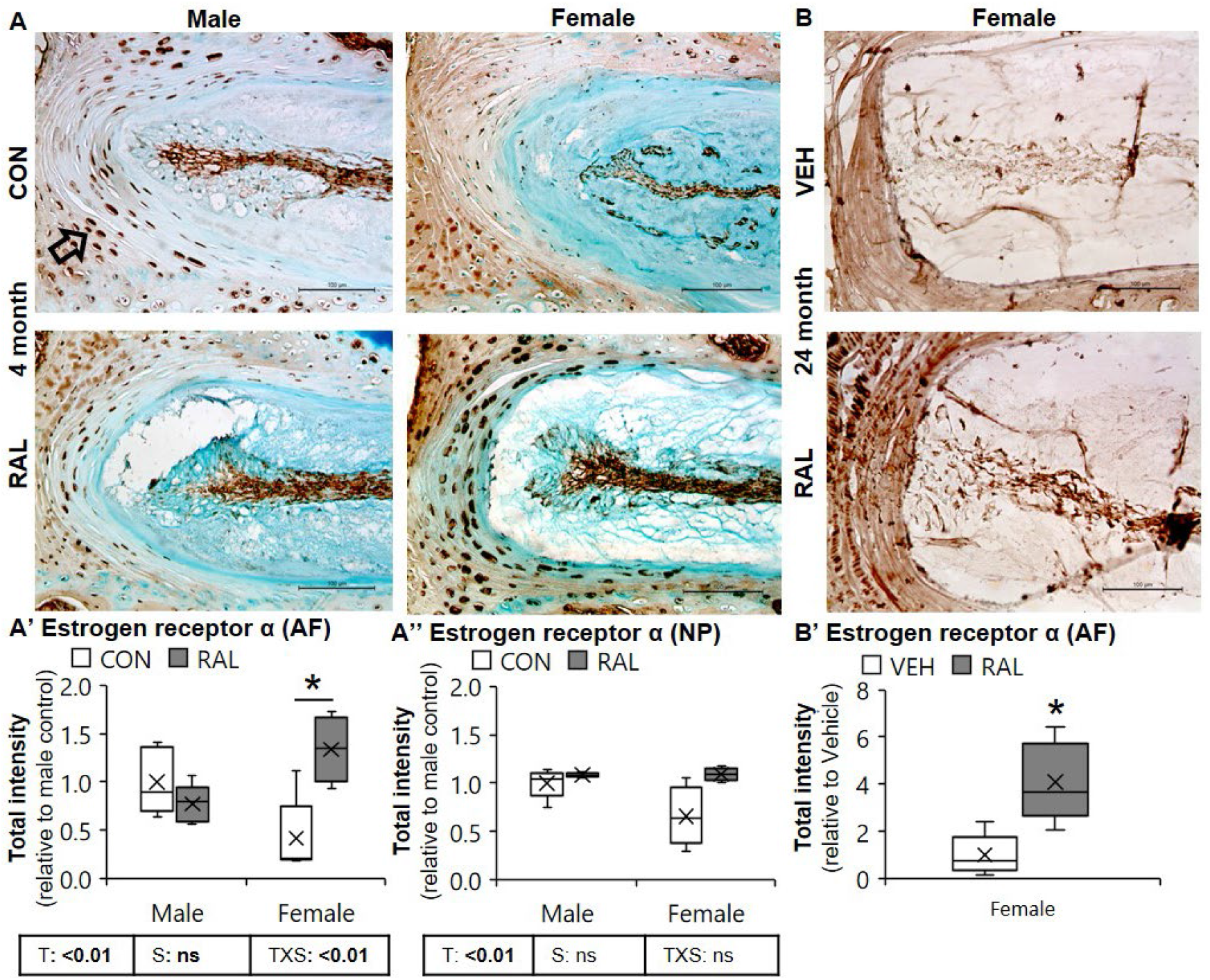
Raloxifene treatment increased the er-α protein expression in female IVDs. cells. (A) 20X representative immunohistochemical estrogen receptor α (er-α) staining. In young IVD, raloxifene increased the er-α protein expression (arrow) in (A’) AF in female IVDs and (A’’) no change in NP in male and female mice. (B) In old IVD, raloxifene increased the er-α protein expression in (B’) AF in female IVDs. Data are represented as box plots with mean marked as cross (x), 25/75% deviation lines and maximum/minimum whiskers. Control (CON, n=5/sex/group) vs Raloxifene (RAL), T: CON vs RAL, S: Male vs Female, TxS: Interaction, Vehicle (VEH, n=8/group) vs RAL, *: VEH vs RAL (female), p<0.05. AF: annulus fibrosus, NP: nucleus pulposus. Scale: 100 μm.

Premenopausal and post-menopausal women experience more back pain compared to age-matched men^(17,18)^. Discogenic pain has long been associated with IVD degeneration^(37)^ and pain-related behavior is noted more so in female mice than males in as early as 12 weeks of age^(6)^. Raloxifene treatment for osteoporosis in women reduces self-assessed back pain^(21)^ and may do so by reducing substance P expression. Here, raloxifene reduced the SP protein expression in the annulus fibrosus by 57% (Fig. 4A, A’) and nucleus pulposus by 38% (Fig. 4A’’) SP-expressing cell numbers followed a similar pattern (Fig. S5). Raloxifene decreased the SP protein expression in AF by 80% in 24-month old IVDs.

**Fig. 4.**
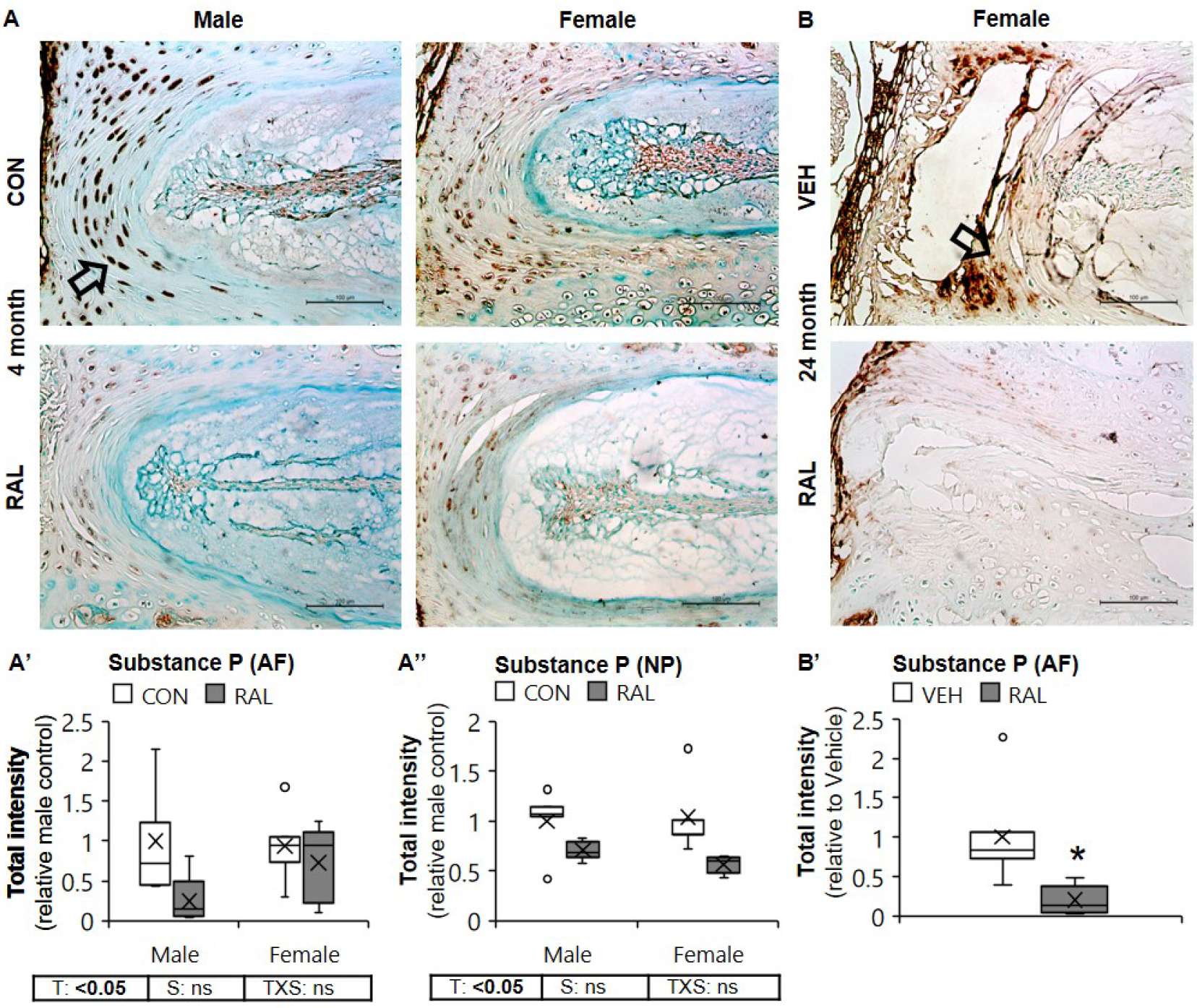
Raloxifene decreased SP expression in NP and AF in young mice and in AF for old female mice. (A) Evaluated using Substance P immunohistochemical staining, raloxifene decreased (A’, A’’) SP protein expression (arrow) in NP and AF in male and female young mice. (B) Evaluated using Substance P immunohistochemical staining, raloxifene decreased (B’) SP protein expression in AF in female old mice. Data are represented as box plots with mean marked as cross (x), 25/75% deviation lines and maximum/minimum whiskers. Control (CON, n=5/sex/group) vs Raloxifene Treated (RAL), T: CON vs RAL, S: Male vs Female, TxS: Interaction, Vehicle (VEH, n=5/group) vs Raloxifene Treated (RAL), *: VEH vs RAL (female), p<0.05. AF: annulus fibrosus, NP: nucleus pulposus. Scale: 100 μm.

### Raloxifene-treated mice developed fluid-filled pockets in the nucleus pulposus of the tail

While there were no significant differences in the tail IVD degeneration score between sexes or treatment with raloxifene in young or old IVD (Fig. S6 A), large ‘pockets’ appeared in young tail IVDs (Fig. 5A). The large pockets were much larger than any one nucleus pulposus cell vacuole and constituted ~9% of the area of the nucleus pulposus in male and female IVDs (Fig. 5A’). To determine whether these pockets contained water, MRI-measured spine segments showed that water content of the tail IVD was not statistically affected by Raloxifene (Table S3). However, raloxifene reduced the incidence of the appearance of the nucleus pulposus cell band (Fig. 5B’). In other words, whereas the cell band (containing no water) was visible in all the control IVDs, the cell band was not visible in 4-out-of-6 raloxifene-treated IVD, indicating presence of water in the band.

**Fig. 5.**
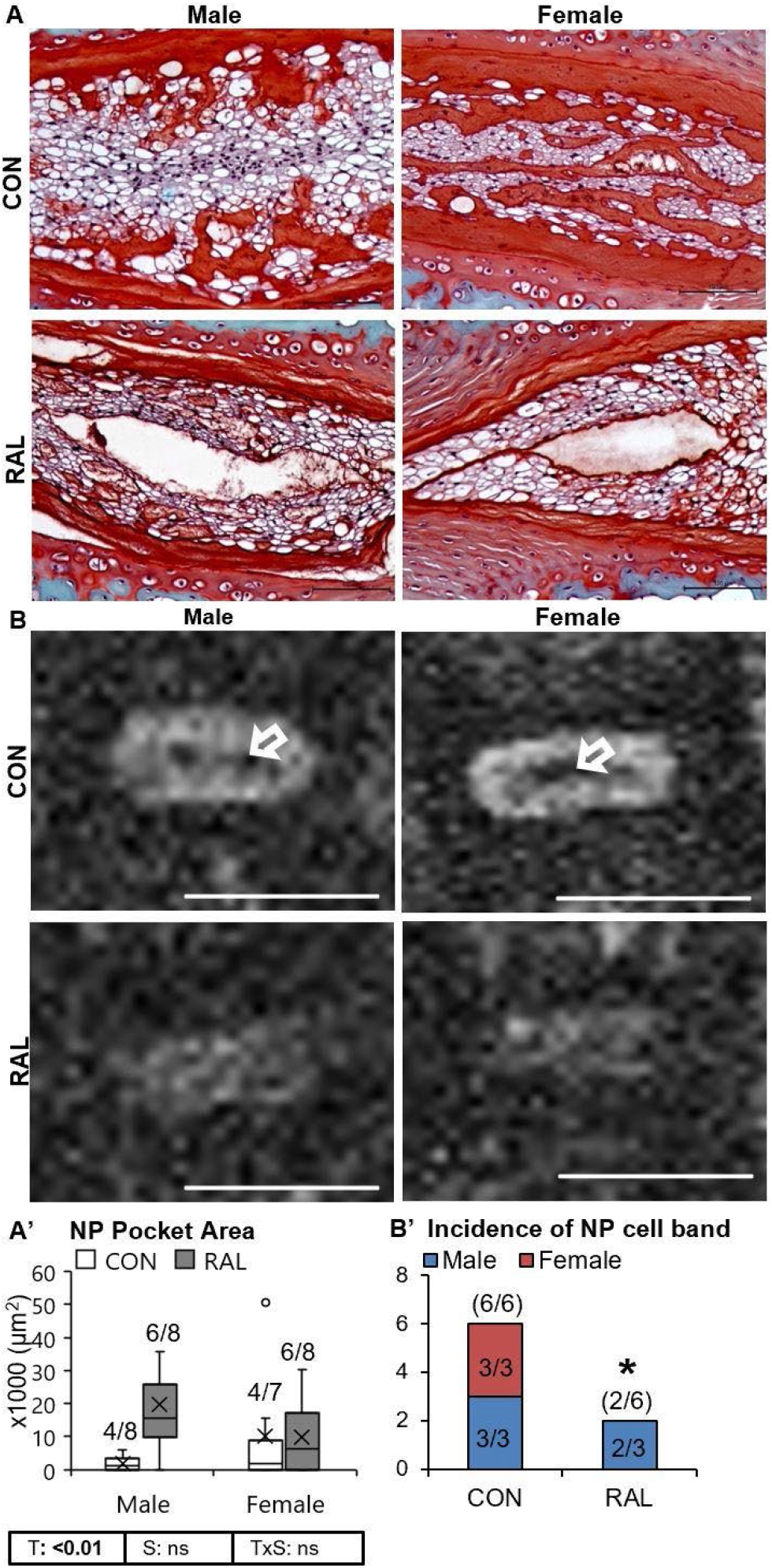
Raloxifene-treated mice developed fluid-filled pockets in the tail nucleus pulposus. (A) Evaluated using Safranin-O/Fast Green staining, (A’) pockets developed in the nucleus pulposus (arrow indicates cell band) (n=7-8/group/sex). (B) Evaluated using MRI, raloxifene (B’) decreased the incidence of absence of fluid in the central nucleus pulposus (n=3/group/sex). Data are represented as box plots with mean marked as cross (x), 25/75% deviation lines and maximum/minimum whiskers. Control (CON) vs Raloxifene (RAL), T: CON vs RAL, S: Male vs Female, TxS: Interaction. *: CON vs RAL, p<0.05. NP: nucleus pulposus. Scale bar: (A) 100 μm, (B) 1 mm.

### Gene expression of tail IVDs were unresponsive to raloxifene

In young tail IVD, raloxifene treatment increased the average gene expression of *er-α* by approximately 2-fold in male and female, with a 10-fold increase in gene expression of male IVDs (Fig. S7A). There were few other changes in gene expression by treatment. Compared to male IVDs, female IVDs had less *col2a1* gene expression (Fig. S7B) and more *lef1* gene expression (Fig. S7E).

### Estrogen-deficiency induces IVD degeneration and increases SP expression in the IVD

To determine whether IVD degeneration and SP are regulated by estrogen, 4-month-old female mice were ovariectomized. Estrogen-deficiency increased IVD degeneration score by 275% (Fig. 6A, A’) and was corroborated by 32% less er-α protein expression in the AF (Fig. 6B, B’). Most of the histological structural changes occurred in the AF, with large spacing and proteoglycan staining between the lamellar fibers, greater AF area and large round inner AF cells (Fig. 6A). In contrast to raloxifene treatment, OVX increased SP protein expression in the AF by 31% (Fig. 6C, C’).

**Fig. 6.**
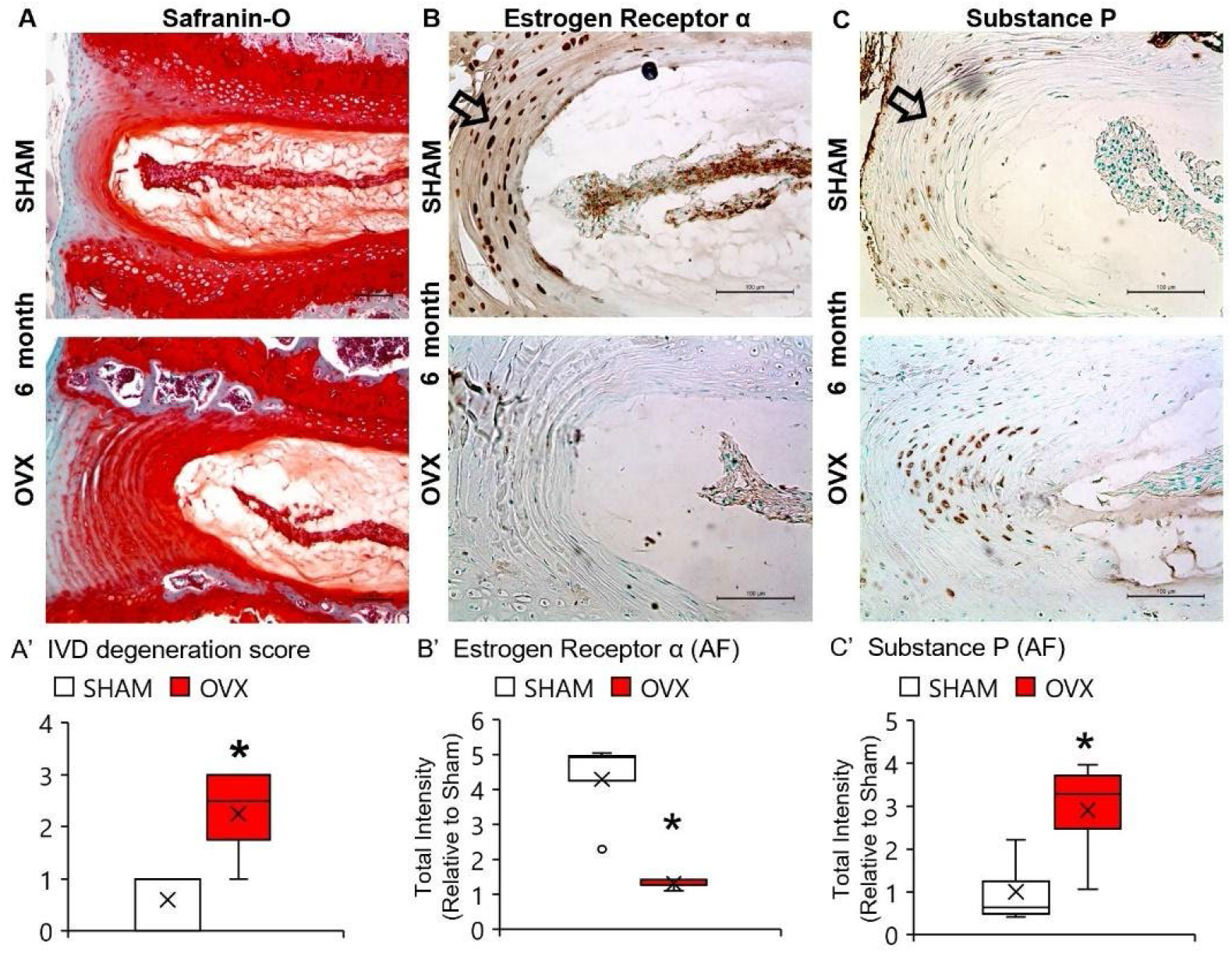
Estrogen-deficiency increased IVD degeneration score and Substance P expression in NP and AF. (A) Evaluated using Safranin-O/Fast Green staining, loss of estrogen increased (A’) IVD degeneration score. (B) Evaluated using er-α staining, (B’) OVX reduced er-α protein expression (arrow) in AF. (C) Evaluated using SP staining, ovariectomy (C’) reduced SP protein expression (arrow) in AF Data are represented as box plots with mean marked as cross (x), 25/75% deviation lines and maximum/minimum whiskers. SHAM (n=4-5/sex/group) vs ovariectomized (OVX), p<0.05. AF: annulus fibrosus, NP: nucleus pulposus. Scale bar: 100 μm.

### By contrast to OVX, raloxifene increased vertebral structural properties and reduced vertebral substance P in osteocytes

We determined the effect of raloxifene on bone structure to corroborate its well-recognized target. In bone, raloxifene increased lumbar trabecular bone volume fraction (BV/TV) by trabecular thickening, with a greater effect in females (Table S2). Tail vertebrae responded similarly. There was no change in cortical bone parameters with treatment (data not shown). With respect to nerve signaling, the number of SP-expressing in female osteocytes was greater compared to that in males, and raloxifene reduced the number of osteocytes that expressed SP by 46% and 55% in males and females young IVDs, respectively (Fig. 7A,A’). Raloxifene treatment reduced the number of SP-expressing osteocytes by 46% in old females. By contrast, OVX increased SP protein expression in vertebral osteocytes (Fig. 8). Lastly, advanced aging increased the number of osteocytes in the control group expressing SP by 46%.

**Fig. 7.**
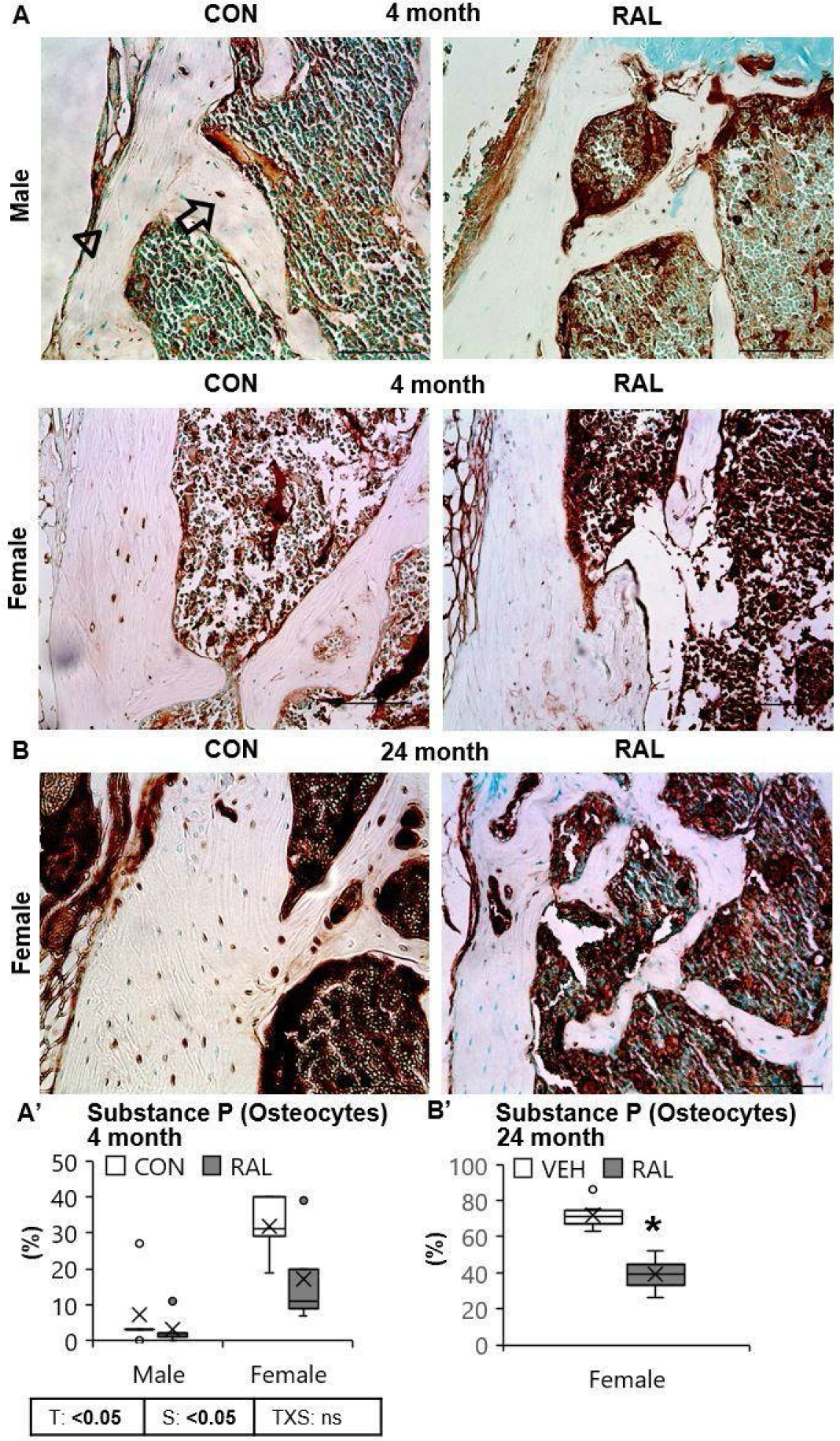
Raloxifene reduced the number of SP-expressing vertebral osteocytes in young and old mice. Estimated in Substance P (arrow vs arrowhead) immunohistochemical images, raloxifene injection decreased the percentage of Substance P expressing osteocytes in L6 vertebra (A, A’) in young male and female mice and, (B, B’) old female mice (n=7-9/sex/group). Data are represented as box plots with mean marked as cross (x), 25/75% deviation lines and maximum/minimum whiskers. Control (CON, n=5/sex/group) vs Raloxifene (RAL), T: CON vs RAL, S: Male vs Female, TxS: Interaction, Vehicle (VEH, n=5/group) vs RAL, *: VEH vs RAL (female), p<0.05. AF: annulus fibrosus, NP: nucleus pulposus. Scale bar: 100 μm.

**Fig. 8.**
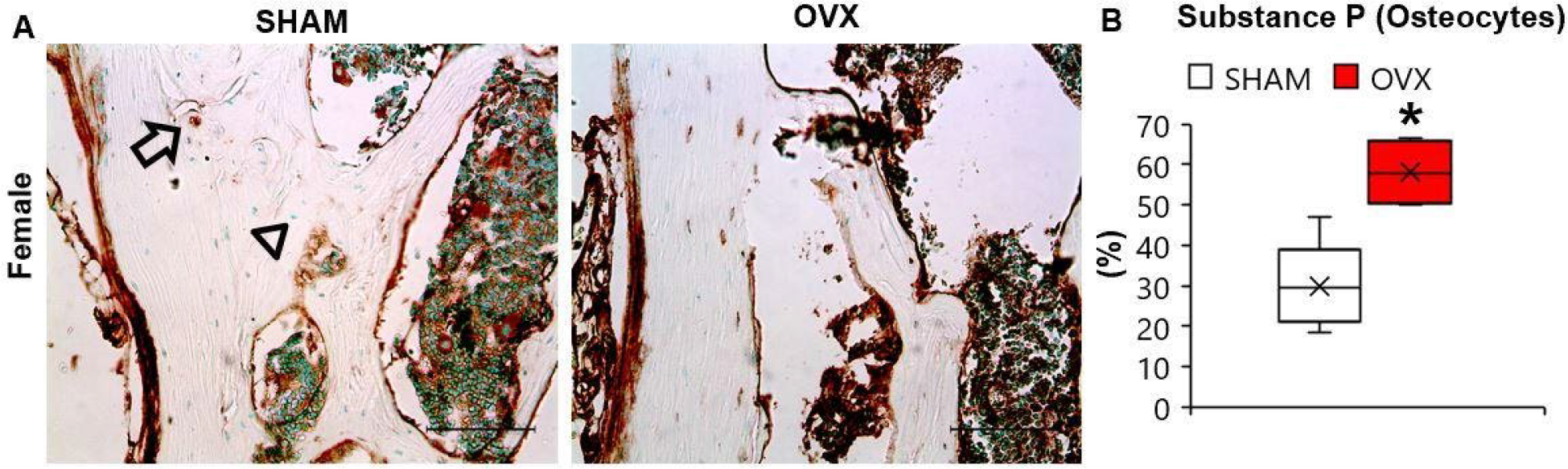
Estrogen-deficiency increased the number of SP-expressing osteocytes in vertebrae. (A) Representative images of 20x Immunohistochemical staining. Ovariectomy led (B) increased the number of SP-expressing osteocytes (arrow vs arrowhead). Data are represented as box plots with mean marked as cross (x), 25/75% deviation lines and maximum/minimum whiskers. SHAM (n=4-5/sex/group) vs ovariectomized (OVX), p<0.05. Scale bar: 100 μm.

## Discussion

We determined the extent of FDA-approved raloxifene to invigorate healthy and degenerating IVDs. Subcutaneous injection of raloxifene prevented the development of sex-related and age-related IVD degeneration, reduced the protein expression of substance P in the IVD/bone of young and old mice, and increased young lumbar IVD structural properties in both sexes by promoting extracellular matrix production, notochordal cell proliferation, ER expression and evenly distributing water-binding. Further, we demonstrated that estrogen-deficiency reduced substance P and induced IVD degeneration. In addition to the widely recognized ability of raloxifene to reinforce bone, these data show that systemic administration of raloxifene promotes major features lost with IVD degeneration and shows great promise at reducing low back pain.

Post-menopausal women incur greater IVD degeneration than men^(46)^ and experience pain more frequently and at higher intensity^(19,20)^. However, whether there is a sexual dimorphism of pain-related behavior in mice is less clear. A retrospective study that compiled the physical behavior of 4,554 mice by open-field activity showed no difference between male and female mice^(47)^. By contrast, some studies show that female mice have more activity than males^(48)^, whereas others show that females have more behavioral signs of pain than males^(6)^. We found that the number of osteocytes that expressed SP was greater in females than males, whereas the number of nucleus pulposus cells that expressed SP was less in females than males. These data suggest that the variable pain-related behavior by sex in mice may be associated with differential spine tissue expression of pain-related markers and highlight the importance of researching the entire functional spinal unit, containing vertebrae surrounding the IVD, and both sexes and ages in the interpretation of pain-related behavior.

Raloxifene prevents osteoporosis-related bone fracture in post-menopausal women by suppressing osteoclast resorption via binding to estrogen receptors. We show that raloxifene also has promise as an IVD therapeutic and we suggest that raloxifene may impact the IVD by at least three mechanisms. Firstly, compared to men, women have low discogenic expression of er-α protein, which may limit raloxifene-induced proliferation^(30,34)^ and both lower er-α and aging can increase pain-related SP in the IVD^(38)^. Therefore, raloxifene may reduce back^(21)^ and joint pain^(49)^ in post-menopausal women because of estrogen-induced reduction of SP in the IVD and bone. Secondly, raloxifene may have increased IVD height^(22)^ by promoting the attraction of water to collagen^(26)^. Lastly, raloxifene stimulated Wnt signaling transcription factor β-catenin, which can also induce IVD anabolism^(8)^. The benefits of raloxifene to stimulate estrogen signaling and reduce substance P expression in IVD and bone extends to old IVD. The relationship between estrogen deficiency, IVD health, and back pain was also established using ovariectomized mice. All these mechanisms will require further investigation to accentuate their effect on the IVD.

Despite several beneficial features, our results implied that raloxifene had some surmountable physiological consequences and that our experimental approach may not have exposed the full translational potential of each therapeutic. First, young tail IVD developed water-filled pockets, which blunted all the anabolic gene expression induced by raloxifene, except for the upregulation of *er-α*. These data suggest that the lower mechanical loading environment of tail IVD compared to lumbar IVD limited the homogeneous distribution of raloxifene and its benefits. Although humans lack tails, these data suggest that recipients of raloxifene should avoid chronic spinal unloading. Lumbar IVDs were not tested by MRI but injection of raloxifene increased IVD area in young mice, which suggests greater hydration. However, the decision to administer raloxifene for IVD degeneration and osteoporosis must be weighed against the greater incidence of stroke and venous thromboembolism^(50)^. We also injected raloxifene in old female mice to demonstrate its efficacy in the population most likely to receive it. While raloxifene reversed the lumbar IVD score to levels similar to young female mice, old tail IVD did not display the ‘pockets’ noted in young mice, which may indicate an age-related impairment of drug transport and may corroborate the lack of raloxifene-induced increase in IVD height in old mice.

In conclusion, there is a great need for pharmacological therapies for IVD degeneration and the transition of regenerative therapies from Bench-to-Bedside includes several developmental stages that can take decades. Raloxifene has a couple of key advantages that can potentially expedite its use: (1) it has been extensively screened for human safety and (2) is already applied to improve an influential tissue of the IVD - bone. Our data suggest that this approach could potentially be implemented in older individuals with severely dehydrated ‘black’ IVD, as commonly noted by MRI, to rehydrate the IVD, prevent IVD collapse and limit discogenic pain. Further study will be required to determine whether this therapeutic prevents severe IVD degeneration. Overall, systemic administration of raloxifene in young and old mice promotes IVD health and its implementation may limit the painful consequences of IVD degeneration in our lives.

## Acknowledgments

Funding: Biomedical Research Grant (EC), NIH AR072609 (JMW), Biomechanics and Biomaterials Research Center grant (NH), Start-up (NH). We would like to thank the Indiana Center for Musculoskeletal Health (ICMH) for their support and thank Dr. Blaine Christensen for kindly providing the ovariectomized and sham-operated spines. Author contributions: N.B.: Study design, assays, data analysis, interpretation of results, and manuscript preparation; P.Z.: assays; O.J.: assays and manuscript preparation; A.B.: assays and manuscript preparation; D.M.: interpretation of results, and manuscript preparation; J.W.: interpretation of results, and manuscript preparation; N.H.: Study design, interpretation of results, and manuscript preparation. Data and materials availability: All data associated with this study are present in the paper or Supplementary Materials.

## Supplementary Materials

**Table S1.** Outcomes for each spinal level.

| **Outcome** | **Lumbar** | **Tail** |
| --- | --- | --- |
| MRI | NA | CC6-7 |
| μCT (Vertebra) | L6 | CC7 |
| qPCR (IVD) | L3-5 | CC8-10 |
| Histology | L1-3 | CC10-12 |

NA: Not Applicable.

**Table S2.**
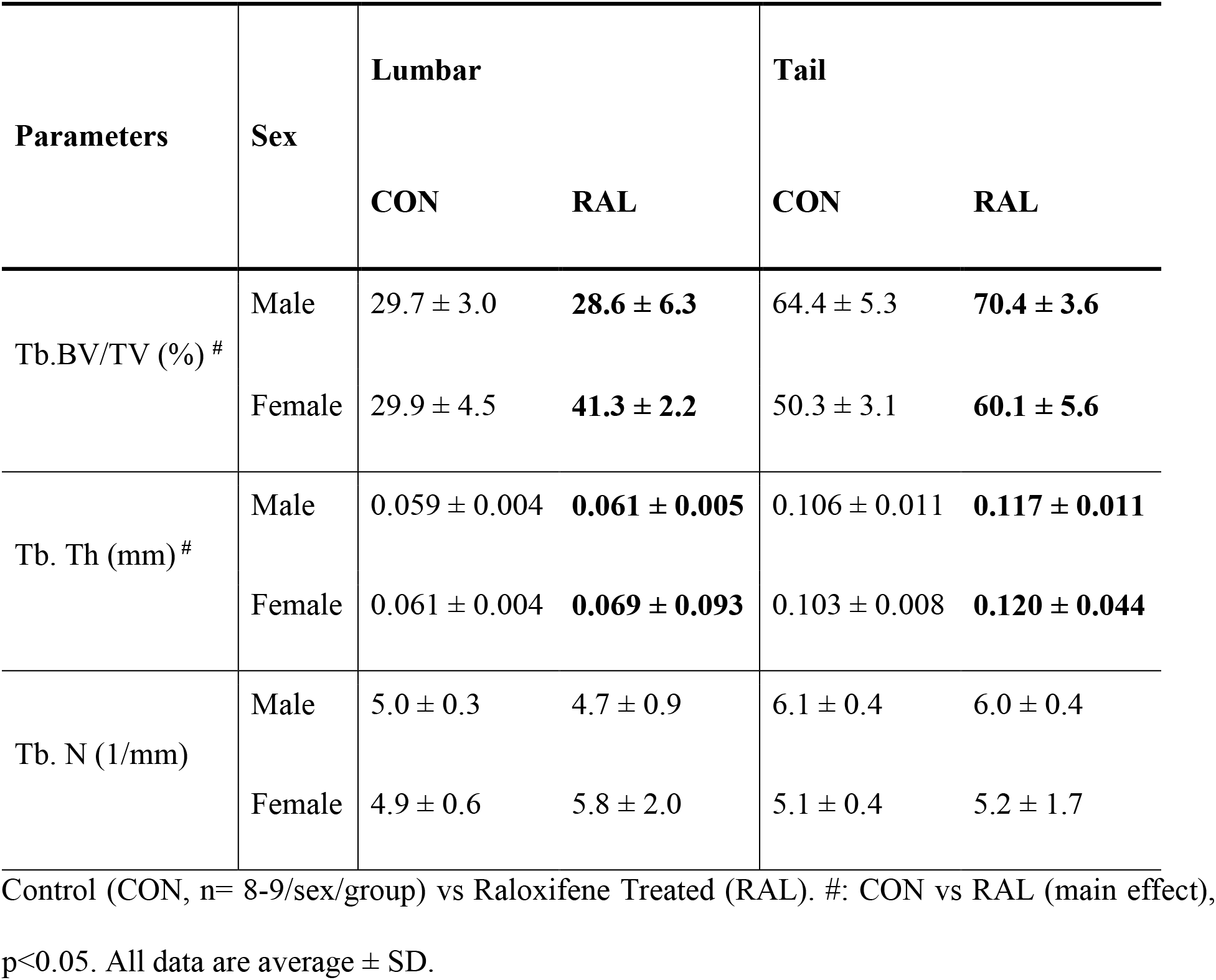
Raloxifene improves lumbar and tail vertebral trabecular structure.

**Table S3.** MRI of spinal segment in mice treated with raloxifene.

| **Sex** | **Group** | **IVD Area (mm<sup>2</sup>)** | **MRI Intensity** | **MRI Index** |
| --- | --- | --- | --- | --- |
| **Male** | **CON** | 0.27 ± 0.19 | 361.61 ± 165.01 | 118.43 ± 116.58 |
|  | **RAL** | 0.32 ± 0.10 | 297.73 ± 121.27 | 99.66 ± 65.43 |
| **Female** | **CON** | 0.30 ± 0.16 | 390.23 ± 94.43 | 124.22 ± 84.32 |
|  | **RAL** | 0.14 ± 0.05 | 270.76 ± 70.75 | 36.41 ± 9.42 |

Using MRI of each animal, IVD area and Intensity were measured using ImageJ and multiplied together to obtain the estimated Water Content value (MRI index). Sample size: Control (CON, 3-4/sex/group) vs Raloxifene Treated (RAL), \*p<0.05.

**Fig. S1.**
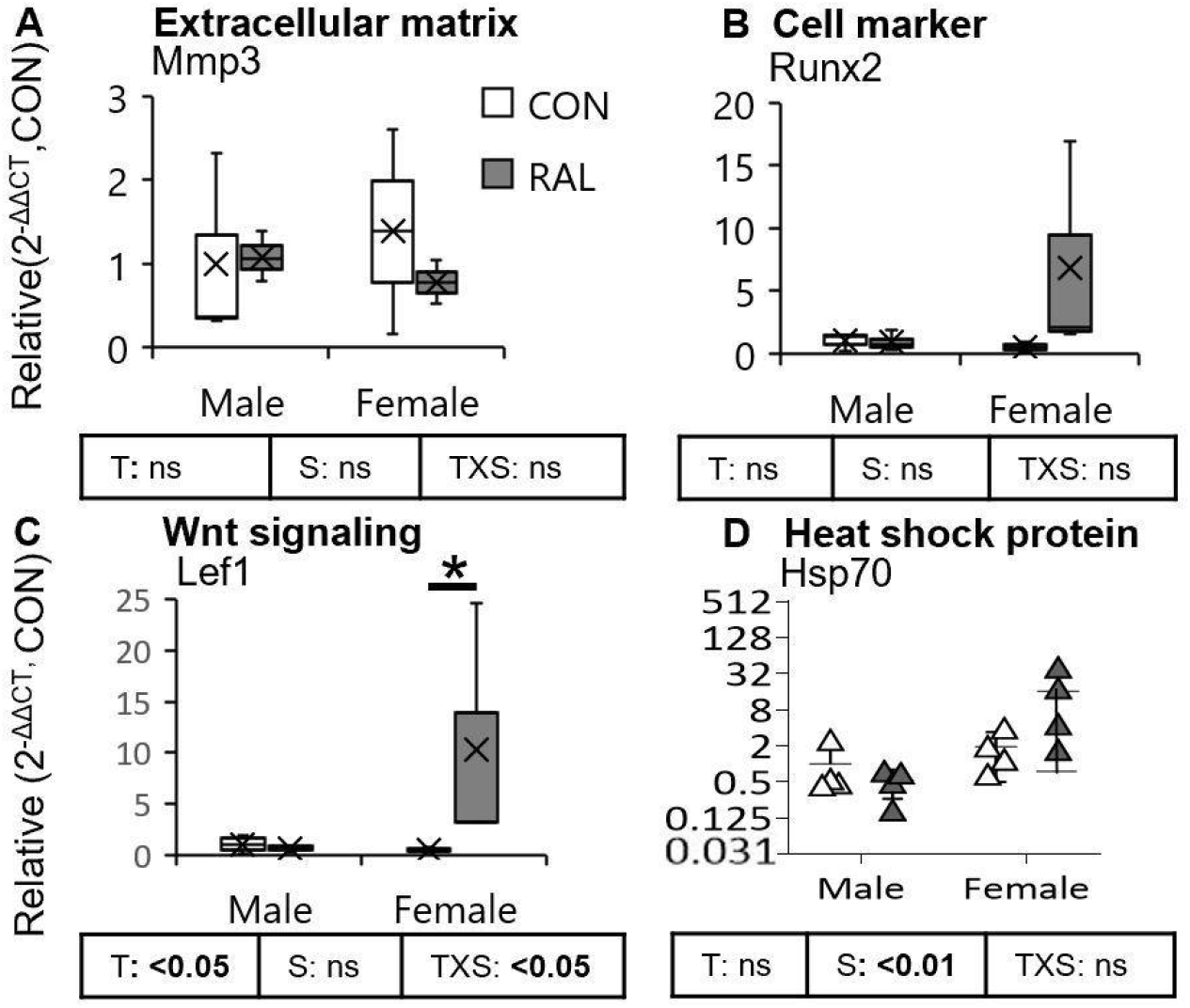
Raloxifene upregulates Wnt signaling in female young mice. Raloxifene treatment (A) did not affect extracellular matrix, *mmp3*, (B) nor chondrogenesis marker, *runx2*, (C) but raloxifene upregulated Wnt Signaling transcription factor *lef1*. (D) Although raloxifene did not affect heat shock protein 70, *hsp70*, female IVD expressed more *hsp70* than male IVDs. Data are represented as box plots with mean marked as cross (x), 25/75% deviation lines and maximum/minimum whiskers or mean ± standard deviation. Control (CON, n=4-5/sex/group) vs Raloxifene Treated (RAL), T: CON vs RAL, S: Male vs Female, TxS: Interaction, *: CON vs RAL (Female). p<0.05.

**Fig. S2.**
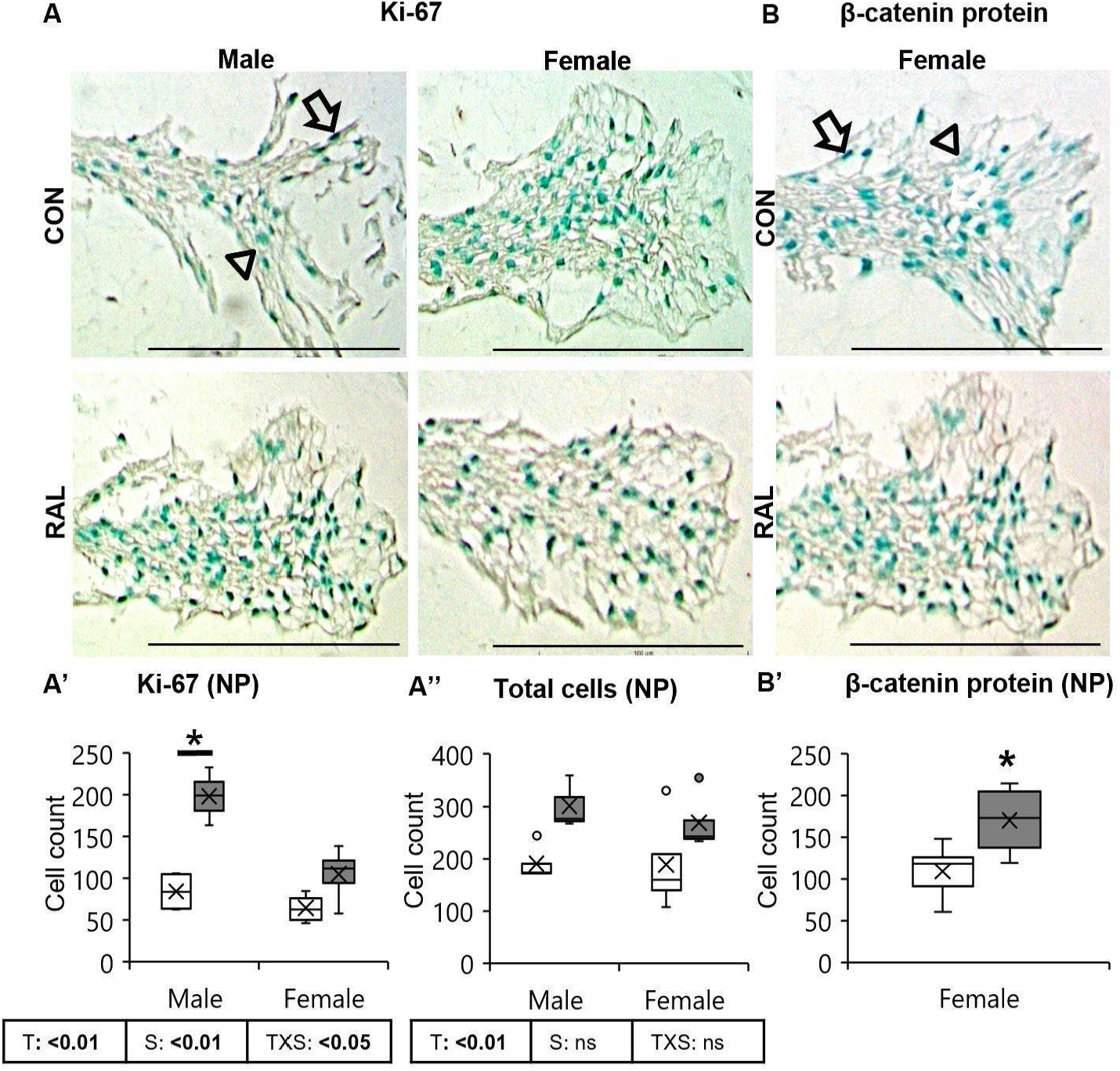
Raloxifene treatment increased the number of ki-67- and β-catenin-expressing cells. (A) Evaluated using images of 20x Ki-67 immunohistochemical staining. Raloxifene increased the (A’) Ki-67 positive cells (arrow) and, (A’’) total cells in the NP. (B) Representative images of β-catenin immunohistochemical staining. (B’) Raloxifene increased β-catenin-positive (arrow) cell count in female IVD. The ROI was a 20x image. (n=4-5/sex/group). Data are represented as box plots with mean marked as cross (x), 25/75% deviation lines and maximum/minimum whiskers. Control (CON) vs Raloxifene-treated (RAL), T: CON vs RAL, S: Male vs Female, TxS: Interaction, *: CON vs RAL (Male/Female),NP: nucleus pulposus, p<0.05. Scale bar: 100 μm.

**Fig. S3.**
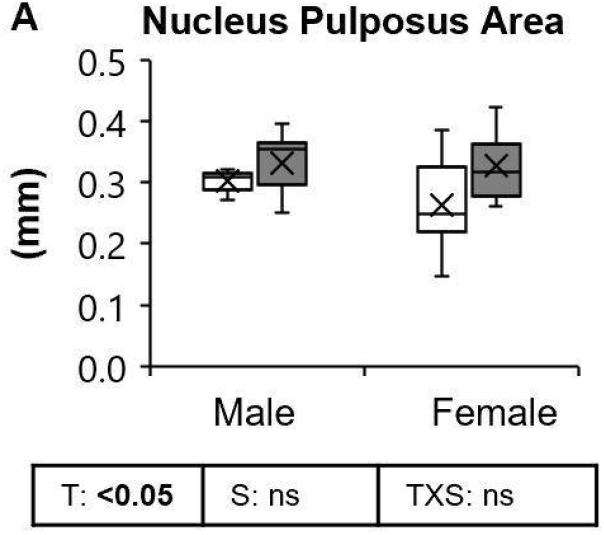
Raloxifene enlarges the nucleus pulposus area in male and female IVDs in mice. Raloxifene (A) increases the NP area of male and female young IVDs. Data are represented as box plots with mean marked as cross (x), 25/75% deviation lines and maximum/minimum whiskers. Control (CON, n=4-5/sex/group) vs Raloxifene Treated (RAL), T: CON vs RAL, S: Male vs Female, TxS: Interaction, p<0.05.

**Fig. S4.**
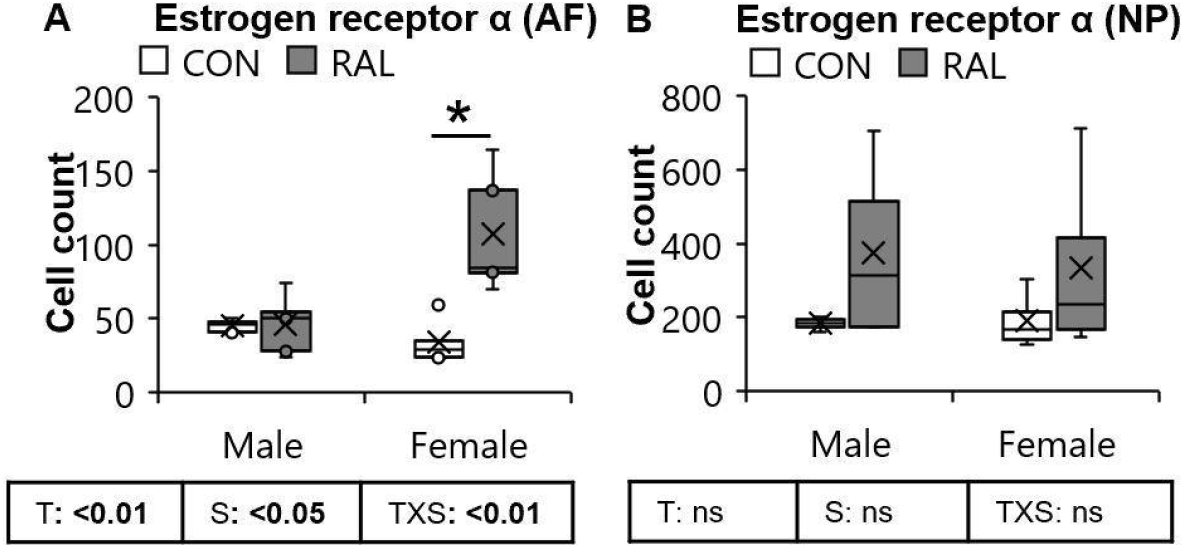
Raloxifene increased the number of er-α protein expressing cells in AF of young female mice. Raloxifene increased the count of Er-α expressing cells in (A) AF and (B) NP in male and female IVD. Data are represented as box plots with mean marked as cross (x), 25/75% deviation lines and maximum/minimum whiskers. Control (CON, n=5/sex/group) vs Raloxifene Treated (RAL), T: CON vs RAL, S: Male vs Female, TxS: Interaction, p<0.05. AF: annulus fibrosus, NP: nucleus pulposus. Scale: 100 μm.

**Fig. S5.**
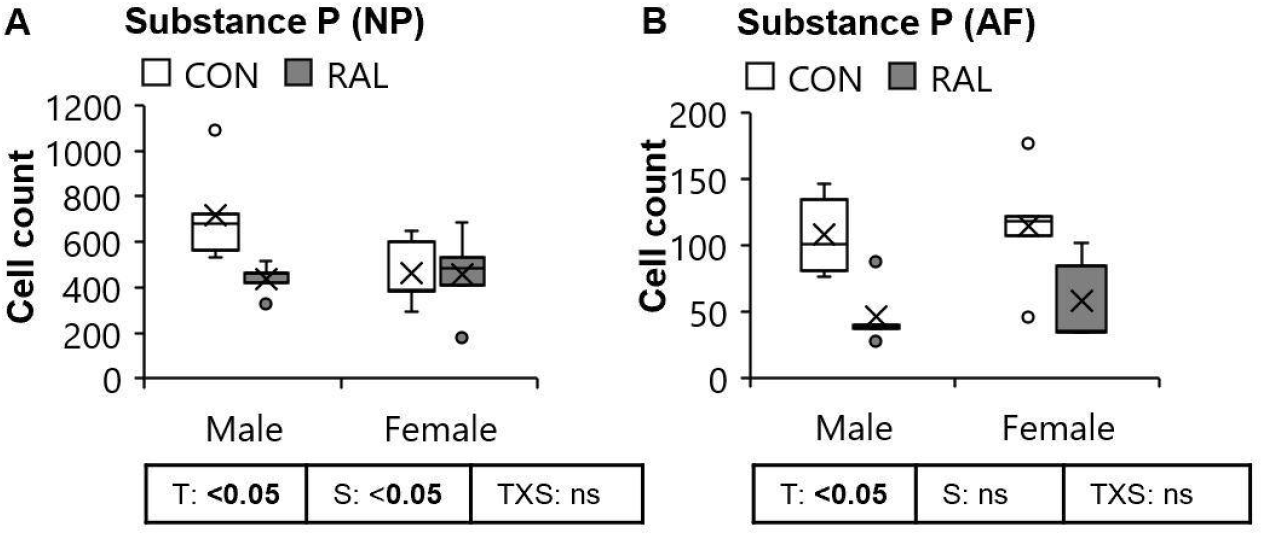
Raloxifene reduced the number of Substance P-expressing cells in male and female IVDs in young mice. Raloxifene treatment reduces the Substance P cell count in (A) NP and, (B) AF of male and female IVDs. Data are represented as box plots with mean marked as cross (x), 25/75% deviation lines and maximum/minimum whiskers. Control (CON, n=4-5/sex/group) vs Raloxifene Treated (RAL), T: CON vs RAL, S: Male vs Female, TxS: Interaction, p<0.05. AF: annulus fibrosus, NP: nucleus pulposus. Scale bar: 100 μm.

**Fig. S6.**
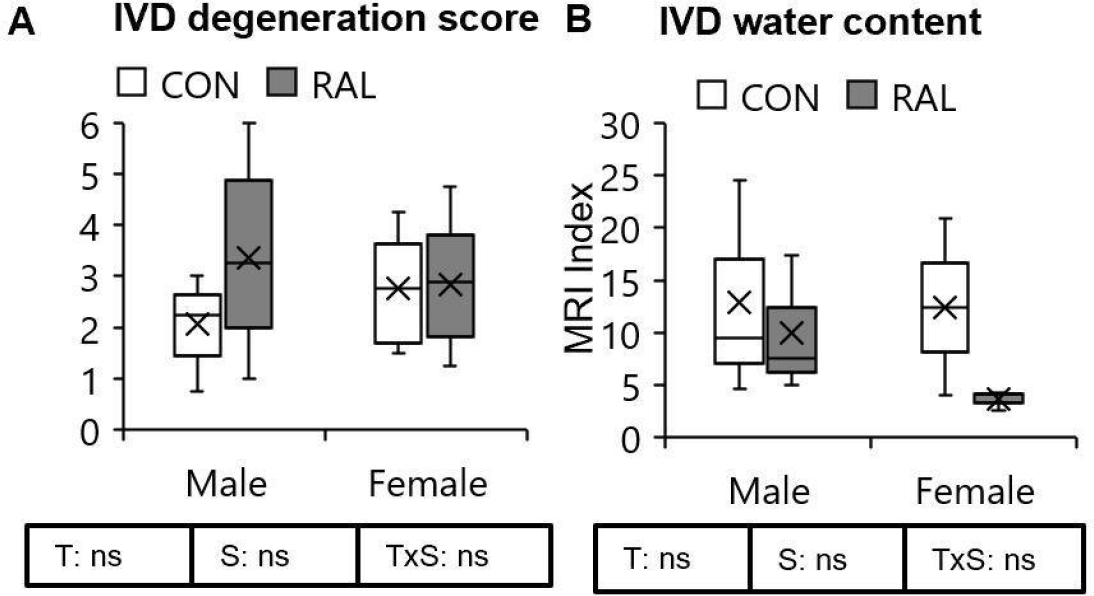
Raloxifene did not improve the IVD degeneration score or water content in tail IVD. Raloxifene had no effect on (A) IVD degeneration nor on (B) water content of tail IVDs. Data are represented as box plots with mean marked as cross (x), 25/75% deviation lines and maximum/minimum whiskers. Control (CON, n=4-5/sex/group) vs Raloxifene Treated (RAL), T: CON vs RAL, S: Male vs Female, TxS: Interaction, p<0.05, Scale bar:100 μm.

**Fig. S7.**
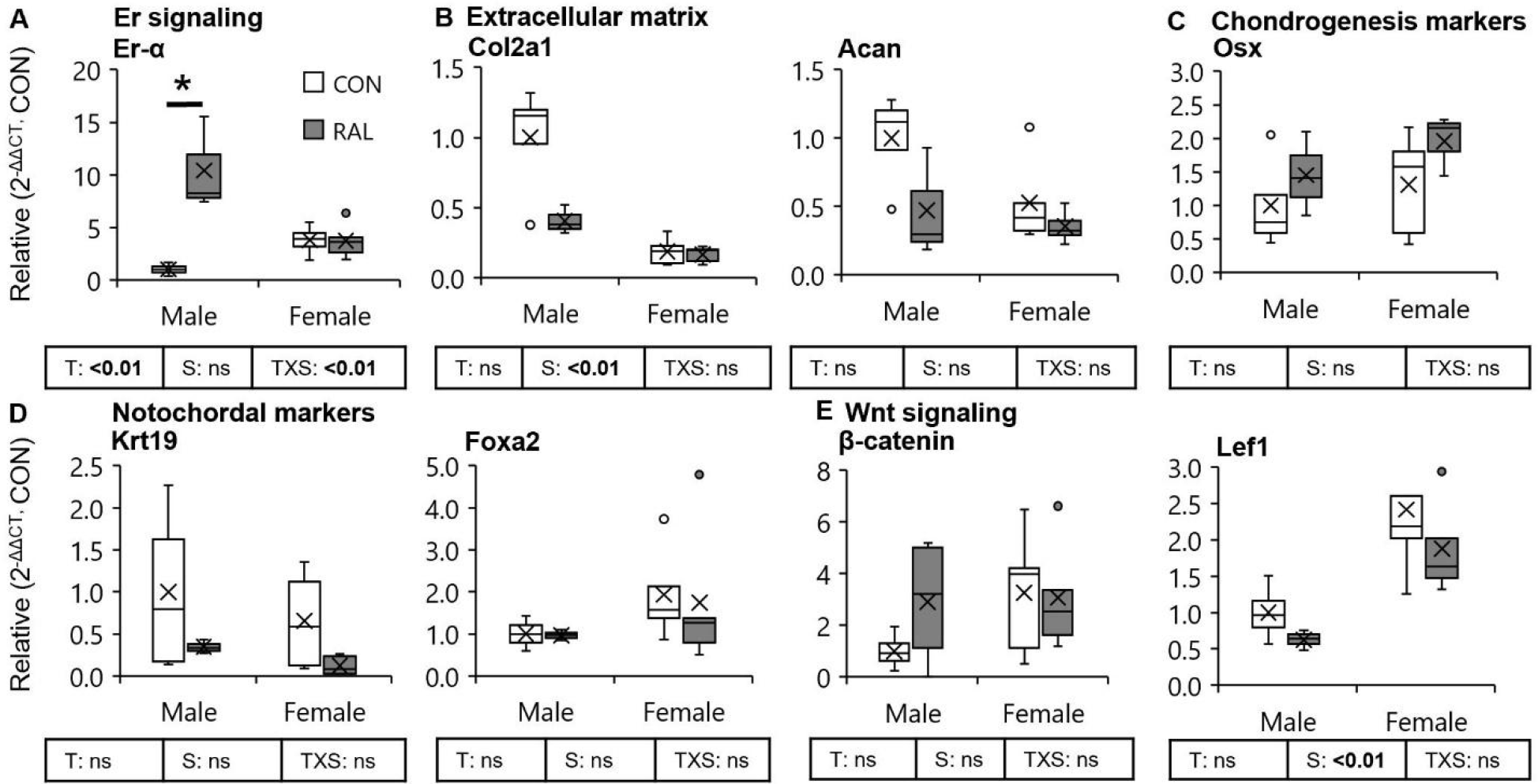
Raloxifene treatment did not alter gene expression in tail IVDs. In tails, other than (A) *er-α* gene expression, (B-E) none of the other gene showed any change with Raloxifene treatment. Data are represented as box plots with mean marked as cross (x), 25/75% deviation lines and maximum/minimum whiskers. Control (CON, n=4/sex/group) vs Raloxifene Treated (RAL), T: CON vs RAL, S: Male vs Female, TxS: Interaction, *: CON vs RAL (Male), p<0.05.

